# A microgel bone marrow model of mesenchymal stem cell paracrine signaling supporting hematopoietic stem cell retention

**DOI:** 10.64898/2025.12.05.692616

**Authors:** Gunnar B. Thompson, Kaila M. Kuo, Andrés J. García, Brendan A.C. Harley

**Author notes:** **Corresponding Author:** B.A.C. Harley, Dept. of Chemical and Biomolecular Engineering, Cancer Center at Illinois, Carl R. Woese Institute for Genomic Biology, University of Illinois at Urbana-Champaign, 110 Roger Adams Laboratory, 600 S. Mathews Ave., Urbana, IL 61801.

## Abstract

Hematopoietic stem cells (HSCs) housed within the bone marrow give rise to the full complement of blood and immune cells. Methods to expand HSCs *ex vivo* have traditionally relied on two-dimensional or liquid culture, but hydrogel approaches have been hypothesized to provide three-dimensional bone marrow-associated biophysical and biomolecular signals that may improve HSC expansion and maintenance *ex vivo*. Here, we describe a granular biomaterial approach to create a multicellular model of the bone marrow. By seeding HSCs amongst mesenchymal stromal cell (MSC)-laden hydrogel microspheres (microgels), we establish paracrine-mediated interactions between HSCs and hydrogel encapsulated MSCs. We provide support for the importance of microgel encapsulation for the emergence of niche-favorable MSC transcriptional profiles. We identify a common cell culture media that accommodates MSC activity while avoiding the use of serum that typically induces differentiation of HSCs. We observe an MSC-density-dependent increase in maintenance of long-term repopulating HSCs in granular co-culture, and we identify significant depletion of long-term repopulating HSCs when both HSCs and MSCs are interstitially seeded in the granular matrix. Together, these findings establish a granular hydrogel co-culture model to examine the influence of MSC-HSC interactions on maintenance and expansion of HSCs in a defined three-dimensional engineered tissue.

## 1. Introduction

Hematopoietic stem cells (HSCs) are the apical cells that sustain the entire hematopoietic system by giving rise to mature, functional blood and immune cells (red blood cells, platelets, granulocytes, lymphoid cells, etc.) through hematopoiesis [1]. These cells primarily reside within the bone marrow of adults in both humans and mice, though HSCs regularly mobilize, trafficking in and out of the bone marrow in small numbers [2]. Hematopoiesis is the differentiation process by which a small number of HSCs that possess multipotency and self-renewal capacity undergo cell division to produce cell populations of progressively reduced stemness to fulfill specific functions such as oxygen transport, fighting infection, and blood clotting. Life-threatening disorders such as leukemia, lymphoma, and sickle cell disease involve dysfunction in the HSC pool [3].

HSC transplants have been administered as a curative therapy for a range of hematopoietic pathologies, but obstacles in donor matching and delivering a sufficient cell dose still limit their application and success. Prior to an allogeneic transplant, the patient must match with a donor, and this is far from guaranteed. In the United States, a patient who requires a transplant has up to an 88% chance of matching if they possess European ancestry, and as low as a 22% chance of matching if the individual is of African descent [4]. Additionally, engraftment success is correlated with the number of total cells [5] and with the number of CD34^+^ (clinical marker for human HSCs) cells delivered [6], and it is not always possible to obtain a sufficient number of cells from the donor. In lieu of an adult donor, it is possible to transplant HSCs from a cord blood unit to restore healthy hematopoiesis in a patient [7, 8]. A benefit to this option is that cord blood transplant presents a lower risk of graft vs. host disease and has less stringent immune matching requirements; however, cord blood units are a less preferred source because they contain less HSCs and present an engraftment failure risk. It has been proposed that methods to expand donor HSCs *ex vivo* could ameliorate limitations regarding the deliverable cell dosage in HSC transplant [8, 9]. Notably, this could also make cord blood a more viable source of HSCs. Because of the reduced immune matching requirements, cord blood expansion may have an extraordinary impact among demographics that historically suffer to find a donor in the United States.

Efforts to expand HSCs often take into account the bone marrow microenvironment, or niche, which provides the combination of mechanical, biochemical, and metabolic cues that regulate hematopoiesis [10, 11]. Niche-associated cues include gradations in stiffness from the endosteum (inner bone surface) to the bone marrow center [12, 13], gradations in oxygen tension from arteriolar to sinusoidal structures [14], as well as complex paracrine cell signaling originating from other niche cells [10]. Many niche cells have been implicated in regulation of hematopoiesis; however, mesenchymal stromal cells (sometimes called mesenchymal stem cells; MSCs) have been observed to provide many of the most critical signals necessary for maintenance of healthy hematopoiesis. Specific subtypes, including Nestin^+^ [15], Lepr^+^ [16], CXCL12-abundant reticular [17], and Ng2^+^ MSCs [18] have been implicated as critical mesenchymal niche cells subtypes (though significant overlap has been noted [10]). These niche cells regulate HSCs through the production of stem cell factor, CXCL12, pleiotrophin, and several other cytokines which are potent effectors of HSC maintenance and retention within the bone marrow.

Researchers have traditionally relied upon animal models that allow for tissue-specific and temporally modulated gene deletion to establish the role of signaling molecules and their cells of origin in HSC regulation. Efforts to expand HSCs have primarily utilized 2D culture methods, and most have aimed to grow HSCs in a medium supplemented with factors including stem cell factor [19, 20], thrombopoietin [21], Flt3 ligand [22, 23], IL-3 and IL-11 [22], or IL-6 [24], which promote stemness and proliferation *in vivo*. These approaches have also included the use of adipose-derived stem cells (ASCs) or MSCs as feeder layers for expansion of (cord-blood derived) HSCs [25, 26]; however in some cases the use of a microporous membrane to separate HSCs form MSCs/ASCs was necessary to increase proliferation and self-renewal [27]. Inspired by the bone marrow niche, efforts have attempted to recapitulate the mechanical and biochemical makeup of the bone marrow using biomaterial systems. Biomaterial approaches, summarized thoroughly elsewhere [28–30], have ranged from marrow-inspired functionalization of polydimethylsiloxane and polyethylene glycol microwells [31, 32] to collagen-coated carboxymethylcellulose [33] or fibronectin-coated polycaprolactone scaffolds [34] that present a 3D macroporous structure. Our lab has previously described the use of gelatin methacrylamide (GelMA) hydrogels that mimic the range of properties present in the bone marrow [35] to study the effect of co-culture of HSCs with MSCs or endothelial cells [12, 36]. Coupled with characterization of diffusive biotransport [37], these approaches provide an avenue to define the relative contribution of paracrine vs. autocrine feedback in a multicellular cohort on HSC fate [38, 39]. We recently developed a maleimide-functionalized gelatin (GelMAL) [40] to enable rapid formation of hydrogel microparticles, or microgels, that enables encapsulation of cells within individual droplets [41]. Microgels present significantly reduced diffusive lengths compared to conventional bulk hydrogels and can be packed to form granular aggregates [42], suggesting avenues to support cell growth and cell-cell communication at scale. We have recently reported rheology-based methods to characterize the yield stress behavior of heterogeneous populations of microgels [43], suggesting routes for *in situ* characterization of an engineered granular bone marrow formed from HSC and MSC-associated microgels. Mixtures of multiple distinct cell-laden microgel populations, or the seeding of discrete cell types in the interstices between cell-laden microgels provides an avenue to present discrete microenvironments to distinct cell populations simultaneously. This strategy suggests ways to effectively shape paracrine interactions while also tuning or eliminating direct cell-cell contact between distinct cell populations.

Here, we describe a microgel-based culture platform to examine the role of MSC-generated paracrine signals on HSC expansion using the well-established murine hematopoiesis model [35]. We benchmark cell activity within GelMAL microgels vs. bulk GelMAL hydrogels (macrogels). We then establish a 3D HSC-MSC microgel culture platform, reporting the effect of the mode of MSC encapsulation (inside vs. between microgels), the weight-percentage of the polymer, and the MSC:HSC ratio on HSC proliferation vs. maintenance. Together, these studies suggest granular assemblies of cell-laden microgels offer the opportunity to create mosaic models of MSC-HSC interactions in the bone marrow that leverage the increased diffusivity of microgel cultures along with the ability to spatially organize distinct cell populations in 3D culture via separate encapsulation in discrete microgels.

## 2. Materials and Methods

### 2.1. Gelatin maleimide synthesis

GelMAL was synthesized as previously described [41]. Gelatin (Type A, Bloom 300, Sigma-Aldrich) was dissolved in 10 mL 2-(*N*-morpholino)ethanesulfonic acid (MES, 1 M, pH 6.0, Gold Biotechnology) and 8 mL dimethyl sulfoxide (VWR) with stirring in a 20 mL vial. After gelatin was fully dissolved, N-hydroxysuccinimide 3-maleimidopropionate (Tokyo Chemical Industry) was dissolved in 2 mL dimethyl sulfoxide and mixed with the gelatin solution. The resultant mixture was adjusted to pH 4.5 using hydrochloric acid and allowed to react for 24 hours at 40 °C with stirring. The reaction products were dialyzed in water acidified to pH 3.25, frozen, and lyophilized to obtain a dry polymer. The degree of functionalization was measured with an ^1^H-NMR spectrometer (Varian Unity Inova 400, 400 MHz). Degree of functionalization was quantified relative to the phenylalanine peak of gelatin. GelMAL used in this study had a degree of functionalization of 0.56 (Fig S1).

### 2.2. Cell culture and medium formulation

C57BL/6 mouse MSCs were purchased from Cyagen (MUBMX-01001) and cultured using Oricell^TM^ complete mesenchymal stem cell growth medium (Cyagen, MUXMX-90011). Cells were received at passage 6 and used at passage 8 – 9, in accordance with manufacturer’s guidelines.

HSC medium was formulated as previously reported [44]: 1X Penicillin-streptomycin-glutamine (Gibco), 10 mM 4-(2-hydroxyethyl)-1-piperazineethanesulfonic acid buffer (HEPES, diluted from 1 M, Gibco), 1 mg/mL polyvinyl alcohol (PVA, 87 – 90% hydrolyzed, Sigma-Aldrich), 1X Insulin-transferrin-selenium-ethanolamine (Gibco), 100 ng/mL thrombopoietin (Peprotech), and 10 ng/mL stem cell factor in Ham’s F-12 Nutrient Mix (Gibco). An alternative HSC media (HSC^-PVA^) was made using all of the listed components except for PVA (to accommodate MSC co-culture).

### 2.3. Microgel formation (acellular and cell-laden)

Microgels were formed using a flow-focusing microfluidic device as described previously [41, 43, 45]. Three solutions were initially made: 1) GelMAL hydrogel precursor in phosphate-buffered saline (PBS, Gibco), 2) 3% v/v SPAN80 (Sigma-Aldrich) in light mineral oil (Sigma-Aldrich), and 3) a 30 mg/mL solution of dithiothreitol (DTT) in PBS. A DTT-in-oil emulsion solution was then made: 4) by combining solutions (2) and (3) in a 3000:237 ratio of oil:DTT and emulsifying by sonication for 90 seconds. To form microgels, solutions (1), (2), and (4) were pumped through the microfluidic device at rates of 5 µL/min, 0.5 µL/min, and 30 µL/min respectively. Acellular microgels were collected in PBS, shaken on ice for 20 minutes to allow complete chemical crosslinking, and then washed with 0.1% Tween-20 (Sigma-Aldrich) in PBS to remove oil. Finally, microgels were washed three times in PBS and then measured, plated, or cultured. Microgels used in this study had a diameter of 198.5 ± 18.6 µm (Fig S2).

Cell-laden microgels were formed in a way similar to a previously described method [41]. Cells were first passaged using TryplE (Gibco) and quenched with MSC medium. Then, cells were washed twice with PBS to dilute serum proteins that would otherwise interfere with DTT/maleimide crosslinking. Cells were concentrated and suspended in hydrogel precursor at the desired concentration prior to microgel formation. Cell-laden microgels were collected in MSC medium, washed in 0.1% Tween-20-in-PBS, and then washed three times in the appropriate medium prior to plating or culture. 25 µL of microgels were formed with MSCs at a density of 4 x 10^6^ cells/mL for MSC viability and gene expression experiments, per replicate. For co-culture experiments, MSC-laden microgels were formed with MSCs at a density of 2.4 x 10^5^ – 3.2 x 10^5^ cells/mL, corresponding to 6000 – 8000 MSCs per sample. Cell-laden microgels (as well as acellular microgels) were cultured in fibronectin-coated 24 well plates (#354411 Corning).

### 2.4. Viability and microgel imaging

Viability was assessed using a LIVE/DEAD cell imaging kit (R37601, Invitrogen). Cells were incubated with the live/dead stain mixture for 15 minutes prior to imaging. Images of cells cultured in 2D were acquired using an EVOS M5000 imaging system (Thermo Fisher). Images of acellular and cell-laden microgels were acquired using a DMi8 Yokogawa spinning disk confocal microscope equipped with a Hamamatsu EM-CCD digital camera (Leica Microsystems, Inc., Deerfield, IL). For viability images, z-stacks were acquired with a step size of 7 µm and a total of 20 slices and compressed into maximum intensity projections. To estimate viability, the live and dead maximum intensity projections were binarized using the Otsu algorithm (this algorithm provided an accurate overlap with qualitatively observed staining). Cell viability was estimated as: *Area_Live_* (*Area_Live +_Area_Dead_*) All image handling and analyses were performed using ImageJ/FIJI. Acellular microgels were imaged for diameter measurements using brightfield and measured using ImageJ/FIJI.

### 2.5. Macrogel formation

GelMAL macrogels were formed by pipetting 20 µL of premixed MSC-laden hydrogel precursor into Teflon molds (dia. 5 mm) fixed to glass slides using a previously described approach [40]. Molds were transferred to a container with ice and covered for 20 minutes to allow for physical crosslinking. After 20 minutes, hydrogels were removed from the molds and transferred to 10 mg/mL DTT for chemical crosslinking in a 24 well plate. Macrogels were allowed to chemically crosslink for 20 minutes. Crosslinking medium was aspirated and replaced with PBS and allowed to wash for 5 minutes on a shaker at room temperature. Washes were repeated for a total of 4 washes before finally resuspending in MSC growth medium.

### 2.6. Gene expression

Gene expression was measured using reverse transcription-quantitative polymerase chain reaction (RT-qPCR). RNA was isolated using the RNeasy mini kit (#74134, QIAGEN) and then reverse transcribed to cDNA using a QuantiTect Reverse Transcription Kit (#205313, QIAGEN) and MyCycler Thermal cycler (Bio-Rad Laboratories). Taqman Fast Advanced Master Mix and Taqman probes for genes of interest were purchased from Thermo Fisher (Table S1) [10, 35, 46–51]. After preparing plates, qPCR was performed using a Real Time PCR QuantStudio 7 (Applied Biosystems). Data was processed using the ΔΔC_T_ method. Gene expression was normalized to *Gapdh* expression for genes measured on each day. Two analyses were performed. First, the gene expression data at Days 4 and 7 were computed as fold-change vs. Day 1 gene expression. For example: Day 4 *Col4a1* expression and Day 1 *Col4a1* expression were normalized to Day 4 *Gapdh* expression and Day 1 *Gapdh* expression, respectively to obtain dCt for each. Then, ddCT was calculated as the difference between Day 4 *Col4a1* and Day 1 *Col4a1* and converted to fold-change. [52, 53]. In a second analysis, the expression of each gene (normalized against *Gapdh* for that condition and that day) was evaluated as the fold-change between microgel-encapsulated MSCs and macrogel-encapsulated MSCs.

### 2.7. 2D media screening

MSC viability assessment in response to distinct media formulations was performed in 2D culture, seeding 7350 MSCs per well in a 48 well plate (to approximate the cell/area of conventional T75 flask culture). Cells were grown in MSC growth medium, HSC medium, a 50:50 blend of MSC and HSC media, or HSC^-PVA^ medium (HSC medium without PVA added). Cells were cultured for two days before evaluating viability.

### 2.8. 3D media screening

MSCs were encapsulated in GelMAL microgels and then seeded in a 24 well plate for culture for either 4 or 6 days. For 4-day media screening experiments, MSCs were cultured in either MSC medium or HSC^-PVA^ medium. Medium was refreshed on day 2 and viability was evaluated after 4 days. Alternatively, MSCs were encapsulated within GelMAL microgels and cultured in MSC medium for 2 days before changing the medium to HSC^-PVA^ medium for an additional 4 days. Medium was refreshed 2 days after this change, with MSC viability was evaluated 4 days after changing the medium.

### 2.9. Hematopoietic stem cell isolation and culture

All animal work was performed under approved animal welfare guidelines (Institutional Animal Care and Use Committee #23033, University of Illinois Urbana-Champaign). Hematopoietic stem and progenitor cells (HSPCs) were isolated as previously described from the tibiae and femora of female C57BL/6 mice aged 6 – 8 weeks (The Jackson Laboratory) [38, 41]. Bones were crushed to release bone marrow into an isolation buffer composed of 5% fetal bovine serum (FBS, R&D Systems) and 0.05 mM ethylenediaminetetraacetic acid (EDTA, Invitrogen) in PBS. Red blood cells were lysed using a 1X red blood cell lysis buffer (BioLegend). Samples were enriched for HSPCs using the EasySep^TM^ Mouse Hematopoietic Progenitor Cell Isolation Kit (#19856, Stemcell Technologies). Cells were then stained for a lineage (Lin) cocktail, Sca-1, c-Kit, and DAPI prior to cell sorting to isolate the Lineage^-^Sca-1^+^c-Kit^+^ (LSK) fraction of hematopoietic cells. Antibodies used for isolation staining are summarized in Table 1. Cells were sorted into HSC^-PVA^ medium using a 5-laser Bigfoot Cell Sorter (Invitrogen). After collection, 6000 – 8000 LSKs were seeded over “pre-cultured” acellular or MSC-laden microgels.

**Table 1.** Stains and antibodies used for prospective isolation of LSKs from mouse bone marrow.

| **Antibody** | **Fluorophore** | **Vendor / Catalog Number** | **μL antibody / μL cell solution** |
| --- | --- | --- | --- |
| N/A | DAPI | BD Biosciences 564907 | 5:300 |
| CD5 | FITC | Thermo Fisher 11-0051-82 | 1:100 |
| B220 | FITC | Thermo Fisher 11-0452-82 | 1:100 |
| CD8a | FITC | Thermo Fisher 11-0081-82 | 1:100 |
| Gr-1 | FITC | Thermo Fisher 11-5931-82 | 0.25:100 |
| Ter-119 | FITC | Thermo Fisher 11-5921-82 | 0.5:100 |
| CD11b | FITC | Thermo Fisher 11-0112-82 | 1:100 |
| Sca-1 | PE | Thermo Fisher 12-5981-83 | 1.25:100 |
| c-Kit | APC-780 | Thermo Fisher 47-1172-82 | 0.625:100 |

### 2.10. Phenotypic analysis of hematopoietic cells after culture

HSC-MSC co-cultures were transferred to Eppendorf tubes and the hydrogel phase was degraded in 200 U/mL collagenase type IV (Worthington Biochemical Corporation). Cells were incubated with an Fc receptor blocker, followed by staining HSC markers using antibodies listed in Table 2. Cells were fixed in formalin prior to flow cytometry using a FACSymphony A1 system (BD Biosciences). Full-minus-one controls were used to set gates for analysis. Long-term repopulating HSCs (LTHSCs) were defined as CD150^+^CD48^-^ LSKs, short-term repopulating HSCs (STHSCs) were defined as CD150^-^CD48^-^ LSKs, and multipotent progenitors (MPPs) were defined as CD150^+/-^CD48^+^ LSKs [22, 54]. Because MSCs were not separated from HSCs in this analysis, MSCs were cultured and subjected to flow cytometry in the absence of HSCs to confirm our ability to continue to use conventional markers to identify murine LSKs. MSCs were negative for lineage markers and c-Kit in flow cytometry experiments, indicating that for HSC analyses, any MSCs would appear within the general cell gate and Lin^-^ gate, but not the LSK gate, and so would not affect the analysis of LSK status or any HSC-specific gates (Fig S3, Fig S4).

**Table 2.** Stains and antibodies used for phenotypic analysis of cultured cells.

| **Antibody** | **Fluorophore** | **Vendor / Catalog Number** | **µL antibody /**<br>**µL cell solution** |
| --- | --- | --- | --- |
| Mouse Lineage Antibody Cocktail | PerCP-Cy5.5 | BD Biosciences 561317 | 20:100 |
| Sca-1 | PE-Cy7 | BD Biosciences 561021 | 0.625:100 |
| c-Kit | R718 | BD Biosciences 567841 | 1.25:100 |
| CD150 | PE | BD Biosciences 562651 | 2.5:100 |
| CD48 | APC | BD Biosciences 562746 | 0.6:100 |

### 2.11. Statistics

Scatterplots and barplots report the mean ± standard error of the mean, unless otherwise noted. The Shapiro-Wilk test was used to test for normality prior to statistical analyses [55, 56], and the Brown-Forsythe test was used to check equivalence of variance between two or more groups [57, 58]. When normality and homoscedasticity were confirmed, conditions were compared using Student’s t-test, One-way ANOVA, or Two-way ANOVA, dependent upon number of groups and experimental design. Tukey’s Honest Significant Differences test was used to compare groups following ANOVA [59]. Welch’s t-test was used when normality was validated, but homoscedasticity was not between two groups [60]. The Kruskal-Wallis test was used to compare groups when normality or homoscedasticity were not valid assumptions [61], and groups were then compared using a Pairwise Wilcoxon Rank Sum test [62]. Plots were generated and statistical tests were performed in R statistical software.

## 3. Results

### 3.1. Mesenchymal stromal cell encapsulation inside of gelatin microgels alters patterns of gene expression

Because of potential exposure to increased shear during microgel encapsulation, we first assessed MSC viability and morphology within 4 wt% GelMAL microgels. MSCs displayed 89.3% ± 0.5% viability and a rounded morphology immediately after encapsulation (Fig 1A-B). By day 1 some MSCs had begun to spread, while by day 3 most MSCs displayed a spread morphology, increased localization toward the microgel surface, and increased density that suggested proliferation. Within 7 days of culture, MSCs fully colonized the surface of the microgels. While viability remained high throughout the 7-day culture, increased fractions of dead cells were observed by day 7 (cell viability: 83.0% ± 7.0%), and many of the dead cells were observed near the center of microgels (Fig S5).

**Figure 1.**
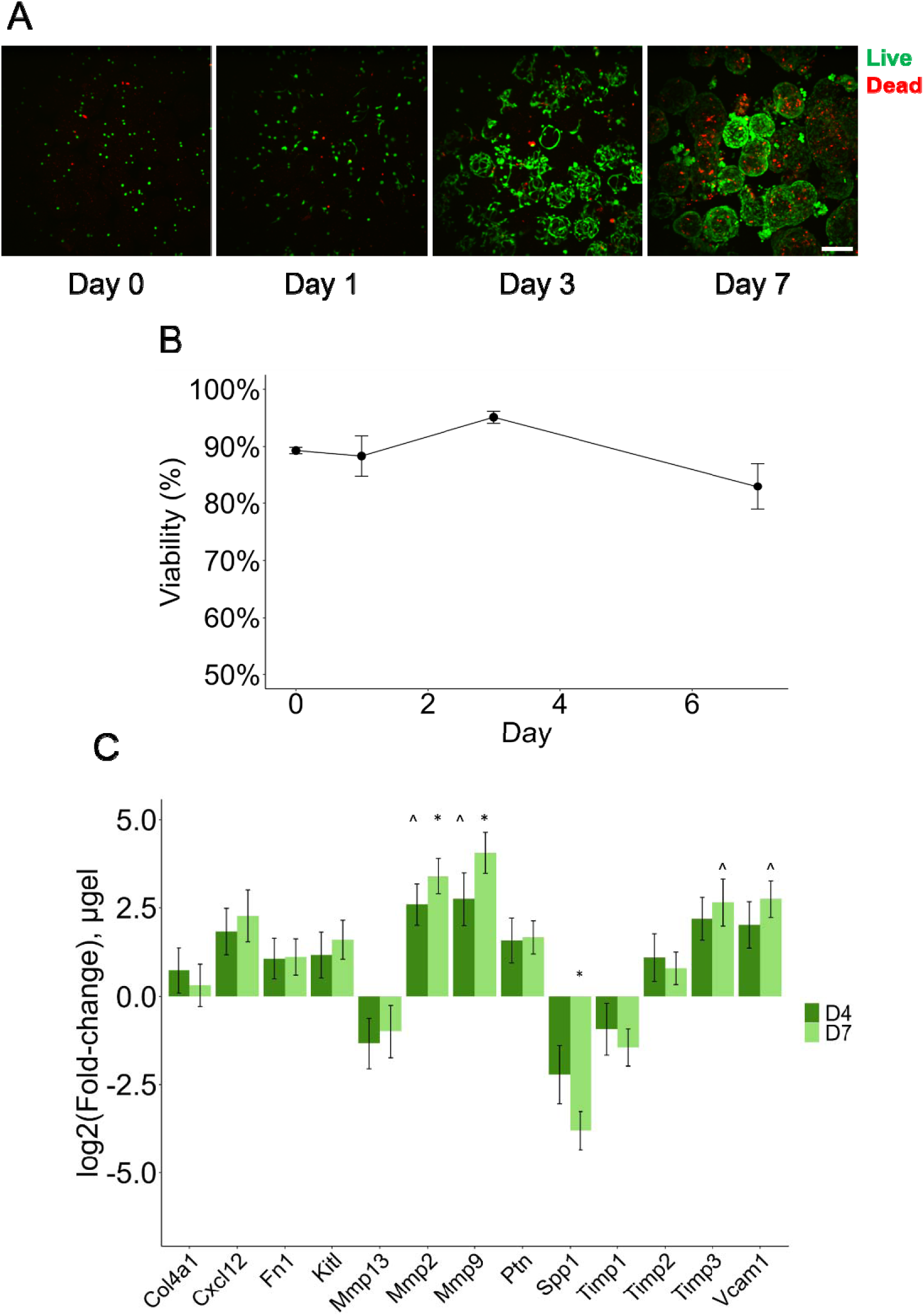
**A** MSCs were encapsulated in 4 wt% GelMAL microgels and cultured for a period of 7 days to assess viability. Scale bar 200 µm. **B** Quantified viability shows that MSCs encapsulated within 4 wt% GelMAL microgels maintain high viability over a period of 7 days. **C** Patterns of gene expression for key remodeling and HSC-regulating genes at days 4 and 7. Data were first normalized to *Gapdh* on the same day, and then fold-change was calculated by comparing against the expression of the gene of interest at day 1. n = 3 independent replicates for each day. ^ indicates p < 0.1 vs. D1. * indicates p < 0.05 vs D1 (t-test).

We then defined the role of microgel vs. macrogel encapsulation on MSC gene expression for a panel of marrow remodeling-associated genes (*Mmp2*, *Mmp9*, *Mmp13*, *Timp1*, *Timp2*, *Timp3*), matrix synthesis genes (*Col1a1*, *Col4a1*, *Fn1*), and HSC-supportive genes (*Kitl*, *Cxcl12*, *Vcam1*, *Spp1*, and *Ptn*). Gene expression data were collected from MSCs encapsulated in 4 wt% GelMAL hydrogels at 1, 4, and 7 days after encapsulation. Expression of remodeling-associated, extracellular matrix (ECM), and HSC-supportive genes trended upward over time for microgel-encapsulated MSCs with only a few exceptions (decreased expression of *Mmp13*, *Spp1*, and *Timp1*; Fig 1C). *Mmp2* and *Mmp9*, associated with ECM degradation, increased significantly by day 7, which likely facilitated the migration of MSCs to the periphery of microgels. The expression of several HSC-supportive genes, including *Cxcl12*, *Kitl*, *Ptn*, and *Vcam1* also increased with time, including significantly increased expression of *Fn1*, *Cxcl12* and *Vcam1* (associated with HSC retention), as well as *Cxcl12*, *Kitl*, and *Ptn* (HSC maintenance) in microgel (vs. macrogel) culture. The expression of *Timp2* and *Timp3* also increased by day 7, suggesting a slowdown in hydrogel matrix degradation. We also compared gene expression for MSCs encapsulated in microgels vs. macrogels (Fig 2). At Day 1, we observed no differences between treatments. Over time, the expression of most genes by microgel-encapsulated MSCs eclipsed macrogel-encapsulated MSCs. By Day 7, the fold-difference between microgel- and macrogel-encapsulated MSCs had become statistically significant for all genes except *Mmp13, Mmp2, Spp1,* and *Timp1*. These data suggest that microgel encapsulation may profoundly improve the gene expression profile of 3D-encapsulated MSCs in the context of bone marrow remodeling and HSC maintenance.

**Figure 2.**
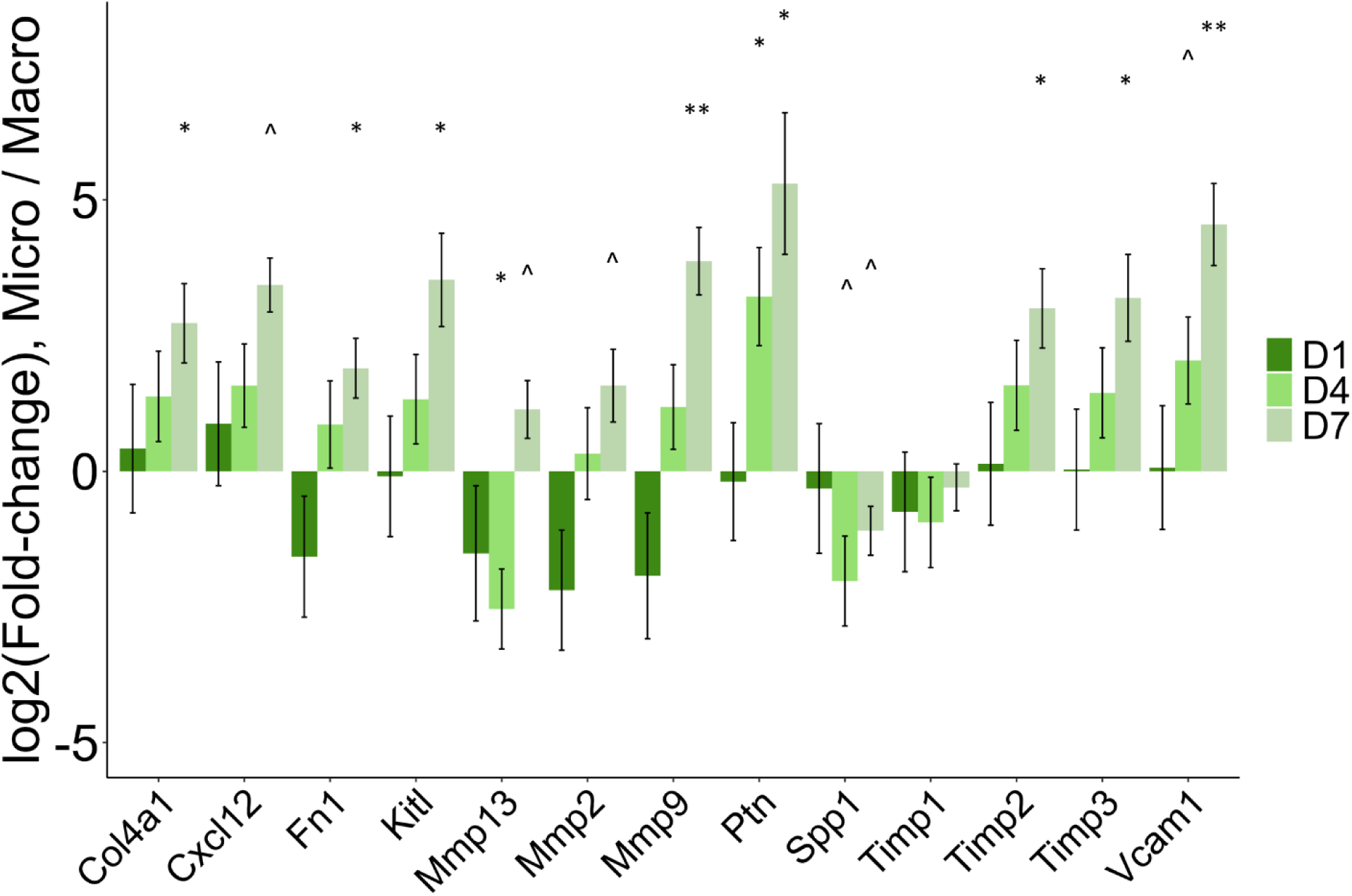
Comparison of gene expression for microgel-encapsulated and macrogel-encapsulated MSCs. For each encapsulation condition and each timepoint, gene expression was normalized to *Gapdh* expression, by timepoint and condition. Fold-change was calculated between microgel-encapsulated MSCs and macrogel-encapsulated MSCs at each timepoint, for each gene. n = 3 independent replicates for each day, for each condition. ^ indicates p < 0.1. * indicates p < 0.05. ** indicates p < 0.01.

### 3.2. Media screening to identify appropriate conditions for hematopoietic and mesenchymal cell co-culture

Direct co-culture of HSCs and MSCs requires a single media formulation to support both cells. We compared a defined HSC expansion medium, identified by Wilkinson et al. [21] and conventional MSC growth medium (that contains undefined factors and FBS). As HSC function is the primary focus of this work, we identified a medium formulation that would best accommodate HSCs in culture and would be tolerated by MSCs. MSC viability was defined after 2 days of culture in MSC medium, HSC medium, or a 50:50 blend of MSC and HSC media (Fig 3A). MSC viability was prohibitively low in HSC medium and reduced in 50:50 MSC:HSC media blends. To better accommodate MSCs in an HSC medium that did not contain any MSC media associated serum, we cultured MSCs in the HSC media containing all components except PVA (designated as HSC^-PVA^). MSC viability was high and growth was similar to the HSC:MSC media blend. As a result, HSC^-PVA^ medium was subsequently used for the rest of the MSC and HSC co-culture experiments.

**Figure 3.**
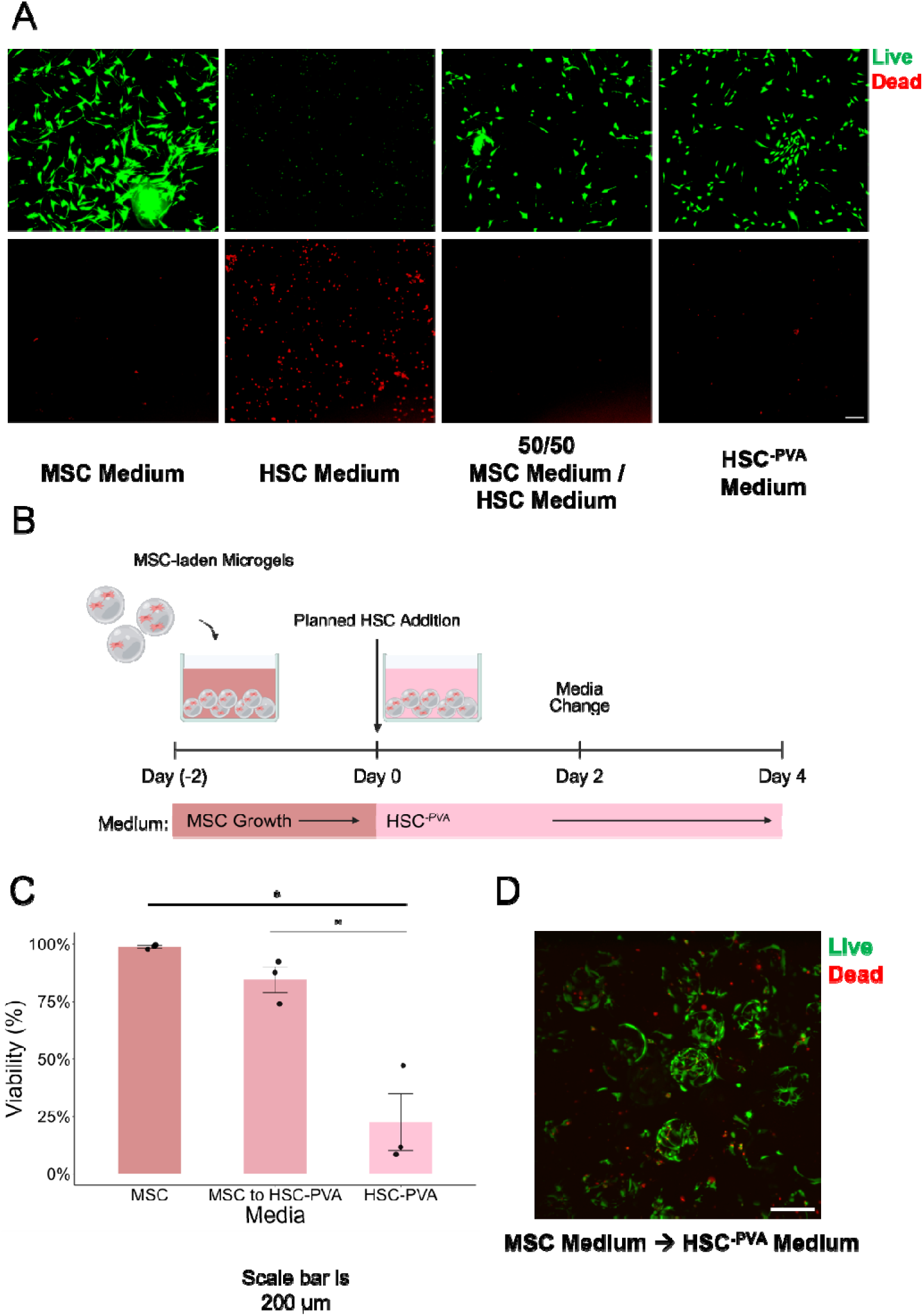
**A** Images show live and dead cell populations for MSCs that were seeded in 2D and cultured in a variety of media. Scale bar 200 µm. **B** Experimental approach for the pre-culture of MSCs with MSC growth medium, followed by a switch to HSC^-PVA^ medium. **C** Viability at the culture endpoint for MSCs encapsulated in 4 wt% GelMAL microgels. When cultured in MSC or HSC^-PVA^ medium only, cells were cultured for 4 days. In the case of pre-culture, MSCs were cultured for a total of 6 days (2 days in MSC medium and 4 days in HSC^-PVA^ medium). * indicates p < 0.05. **D** Representative image of MSCs after pre-culture in MSC medium and 4-day culture in HSC^-PVA^ medium. Scale bar 200 µm.

We then evaluated MSC viability as a function of culture medium when encapsulated in 4 wt% GelMAL microgels for 4 days (the target culture duration for HSCs). As expected, the MSC medium accommodated high viability and growth; however, despite the excellent viability observed when MSCs were cultured in 2D in HSC^-PVA^ medium alone, MSCs cultured in microgels fared poorly in HSC^-PVA^ medium (Fig S6). To address this problem, we adopted a new experimental approach in which MSCs would be encapsulated and initially “pre-cultured” in MSC growth medium for two days with the goal to reduce damage form encapsulation, prior to a medium change to the HSC^-PVA^ medium coinciding with introduction of HSCs for subsequent culture (Fig 3B). When cultured in this manner, MSC viability was much improved, reaching 84.6% ± 5.5% at day 4 of HSC^-PVA^ medium exposure, relative to the HSC^-PVA^ medium with 22.4% ± 12.3% viability (Fig 3C-D). This approach was used for all subsequent HSC-MSC co-culture studies.

### 3.3. Phenotypic analysis of hematopoietic stem cells after co-culture with microgel-encapsulated mesenchymal stromal cells

MSCs were encapsulated in 4 wt% or 6 wt% GelMAL microgels and then seeded in 24-well plates at day −2 in conventional MSC media, with acellular microgels fabricated as a control. 4 wt% is a continuation from previous work [41], and 6 wt% was chosen to present a matrix density similar to our previous work in GelMA that supported HSC maintenance [35]. Media was replaced with HSC^-PVA^, and HSCs were seeded interstitially into packed beds of MSC-laden or acellular granular gels at day 0. Cultures were maintained for 4 days, then collected and processed for flow cytometry at day 4. The wt% of GelMAL had minimal impact on HSC phenotype: no effects were observed in the LTHSC or MPP populations. Co-culture with 6 wt% microgels led to an increase in the number of STHSCs at Day 4 as a factor, but there were no statistical differences observed between individual conditions on the basis of wt % (Fig S7). The overall number of Lin^-^ cells present at the end of culture was elevated at least in part due to MSCs appearing within the Lin^-^ gate (Fig 4A-B). Lin^+^ cell numbers were slightly elevated with MSC co-culture (Fig 4C). The number of retained LSKs increased with MSC co-culture (Fig 4D). When LSKs were co-cultured with MSC-laden microgels, the number of LTHSCs was unaffected, while the number of STHSCs was reduced and the number of MPPs increased (Fig 4E-G). Taken together, these data suggest that MSCs seeded within microgels increase the number of LSKs maintained and promote differentiation of STHSCs toward MPPs, inducing expansion of (more differentiated) Lin^+^ cells population. Interestingly, these downstream progeny changes occurred without losses in the LTHSC population, indicating distinct responses by distinct LSK subpopulations.

**Figure 4.**
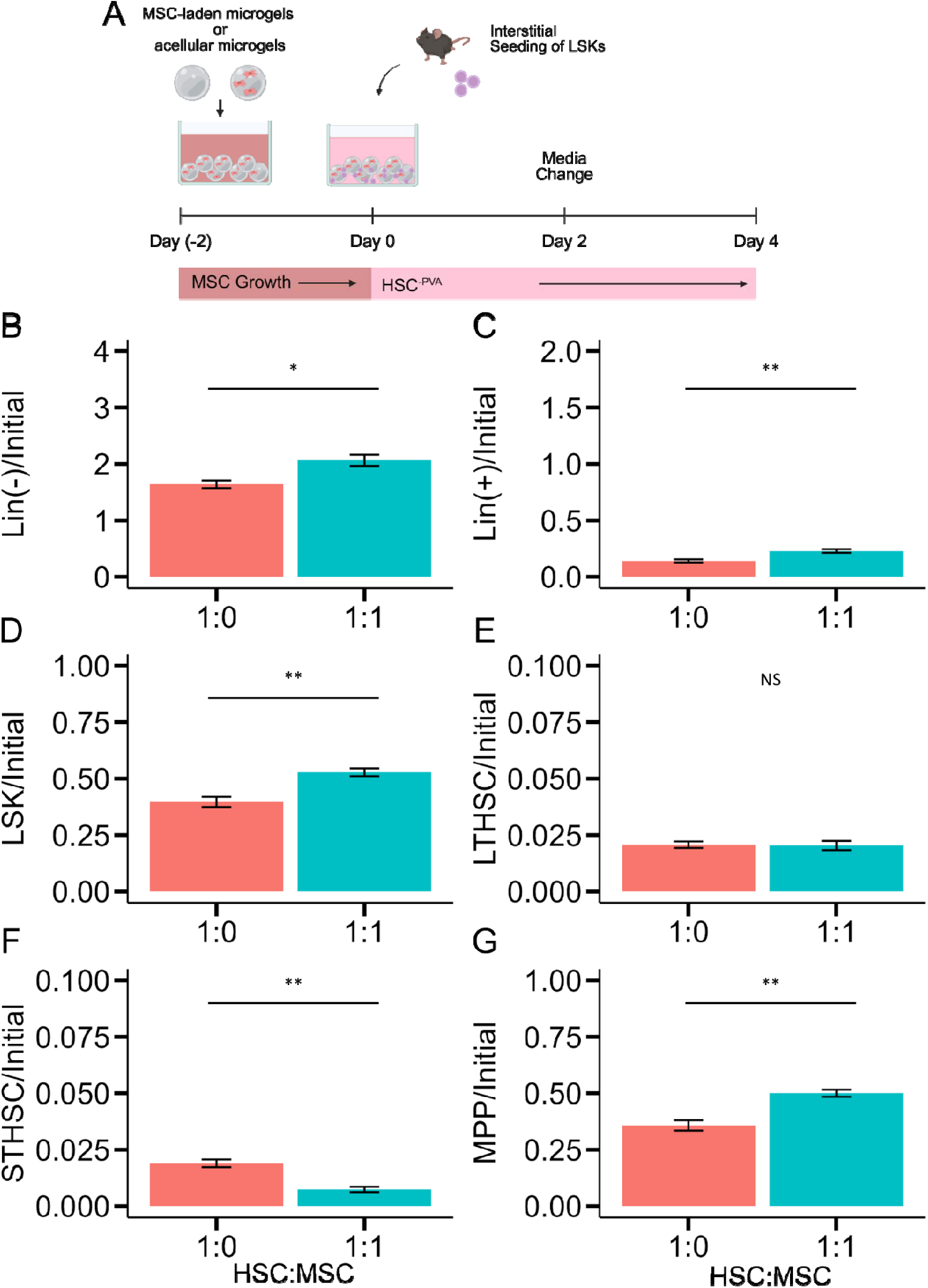
**A** HSCs were interstitially seeded in loosely packed beds of either acellular or MSC-laden 4 wt% GelMAL microgels and cultured for 4 days prior to flow cytometry analysis. MSCs were encapsulated within microgels. Plots present **B** Lin^-^ cells normalized to the initial seeded number of cells, and **C** Lin^+^ cells normalized to the initial seeded number of cells. The HSPC populations were analyzed and normalized against the initial number of ceded cells as follows: **D** LSKs, **E** LTHSCs, **F** STHSCs, **G** MPPs. * p < 0.05, ** p < 0.01.

### 3.4. Increasing the ratio of mesenchymal to hematopoietic cells enhances maintenance of long-term repopulating hematopoietic stem cells

Having seen an effect of MSC co-culture on HSC progeny, we next aimed to better define the relationship between the numbers of MSCs and HSCs in this system by increasing the MSC:HSC ratio and measuring HSC phenotype. We evaluated HSC monoculture (HSC cultures with acellular microgels), 1:1 HSC:MSC co-cultures, and 1:10 HSC:MSC co-cultures by again seeding MSCs at day −2 and adding HSCs at day 0 (Fig 5A). The overall number of Lin^-^ at day 4 were similar for niches containing 1X or 10X MSCs (Fig 5B). Consistent with earlier results, the Lin^+^ fraction of hematopoietic cells increased with MSC density (Fig 5C), whereas the number of LSKs decreased when the ratio of MSC:HSC was increased from 1:1 to 10:1 (Fig 5D). Most notably, increasing the fraction of resident MSCs increased the size of the remaining LTHSC population (Fig 5E), but at the expense of decreased numbers of downstream STHSCs and MPPs (Fig 5F-G). These results suggest that LTHSCs are better maintained in the presence of MSC-laden 4 wt% GelMAL microgels.

**Figure 5.**
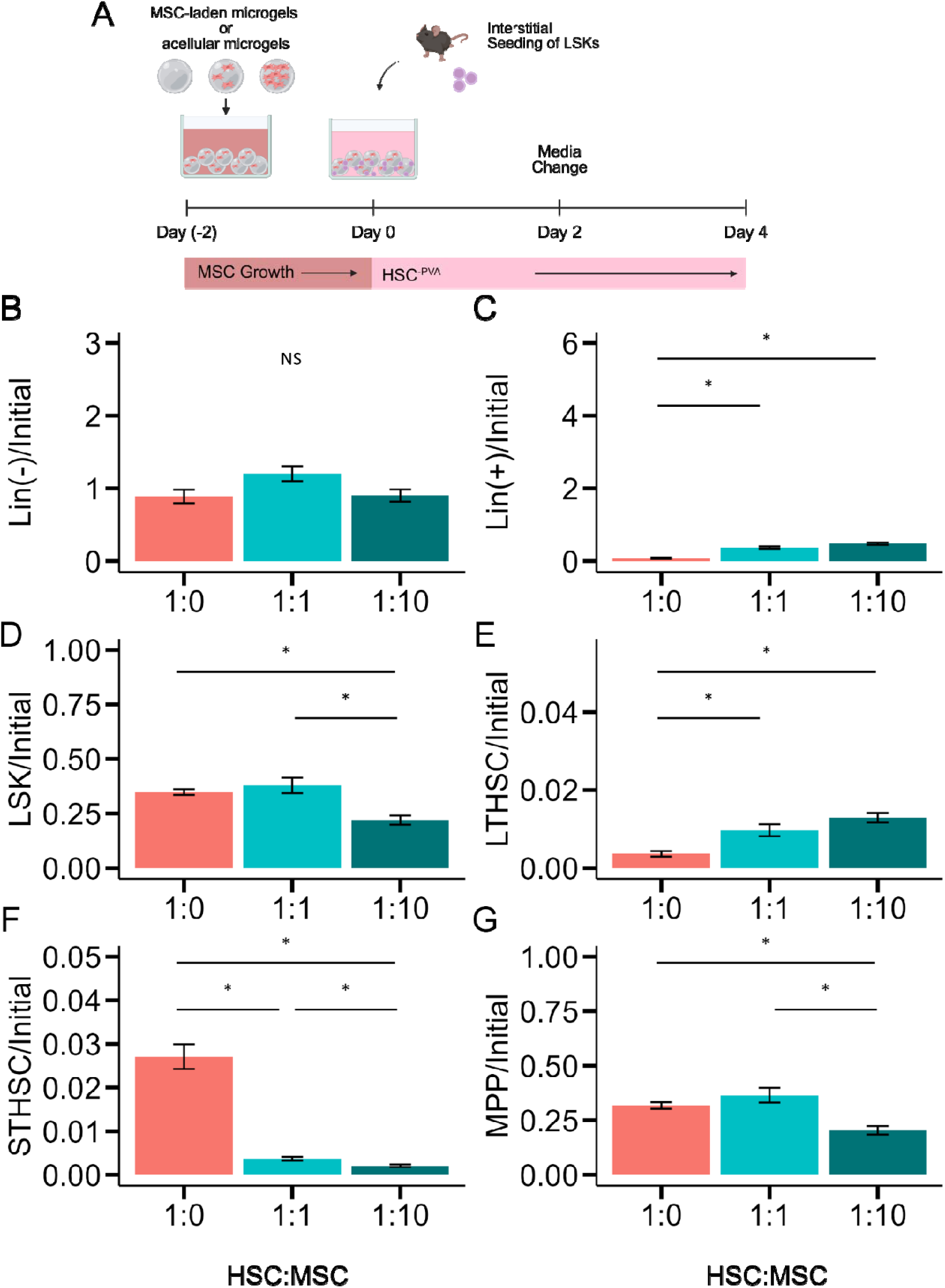
**A** HSCs were interstitially seeded in loosely packed beds of either acellular or MSC-laden microgels and cultured for 4 days prior to flow cytometry analysis. MSCs were encapsulated within microgels. Two ratios of MSC:HSC were tested – 1:1 and 10:1. Plots present **B** Lin^-^ cells normalized to the initial seeded number of cells, and **C** Lin^+^ cells normalized to the initial seeded number of cells. The HSPC populations were analyzed and normalized against the initial number of ceded cells as follows: **D** LSKs, **E** LTHSCs, **F** STHSCs, **G** MPPs. * p < 0.05.

### 3.5. Cell-cell contact in the granular model induced significant differentiation of hematopoietic stem cells

Microgel-based granular assemblies also allow for the facile incorporation of multiple cell populations interstitially. We briefly investigated the effect of seeding both HSC and MSC populations interstitially within the granular model, which allows for a greater degree of direct cell-cell contact between the HSCs and MSCs (Fig 6A). While the Lin^-^ population remained unchanged (Fig 6B), a 4-fold to 5-fold expansion of Lin^+^ cells was observed (Fig 6C), indicating significant hematopoietic differentiation and proliferation. While the number of LSKs was unchanged when moving MSCs from inside microgels to the interstices between microgels with the HSCs (Fig 6D), we observed significant alterations in the composition of the LSK compartment. Notably, LTHSCs and STHSCs were nearly depleted (Fig 6E-F). MPP numbers were unchanged in 1:1 HSC:MSC interstitial co-culture (Fig 6G). Combined with the observation of Lin^+^ expansion, this result suggests a greater amount of differentiation within the MPP subpopulation when HSCs are co-cultured with MSCs interstitially in this granular material.

**Figure 6.**
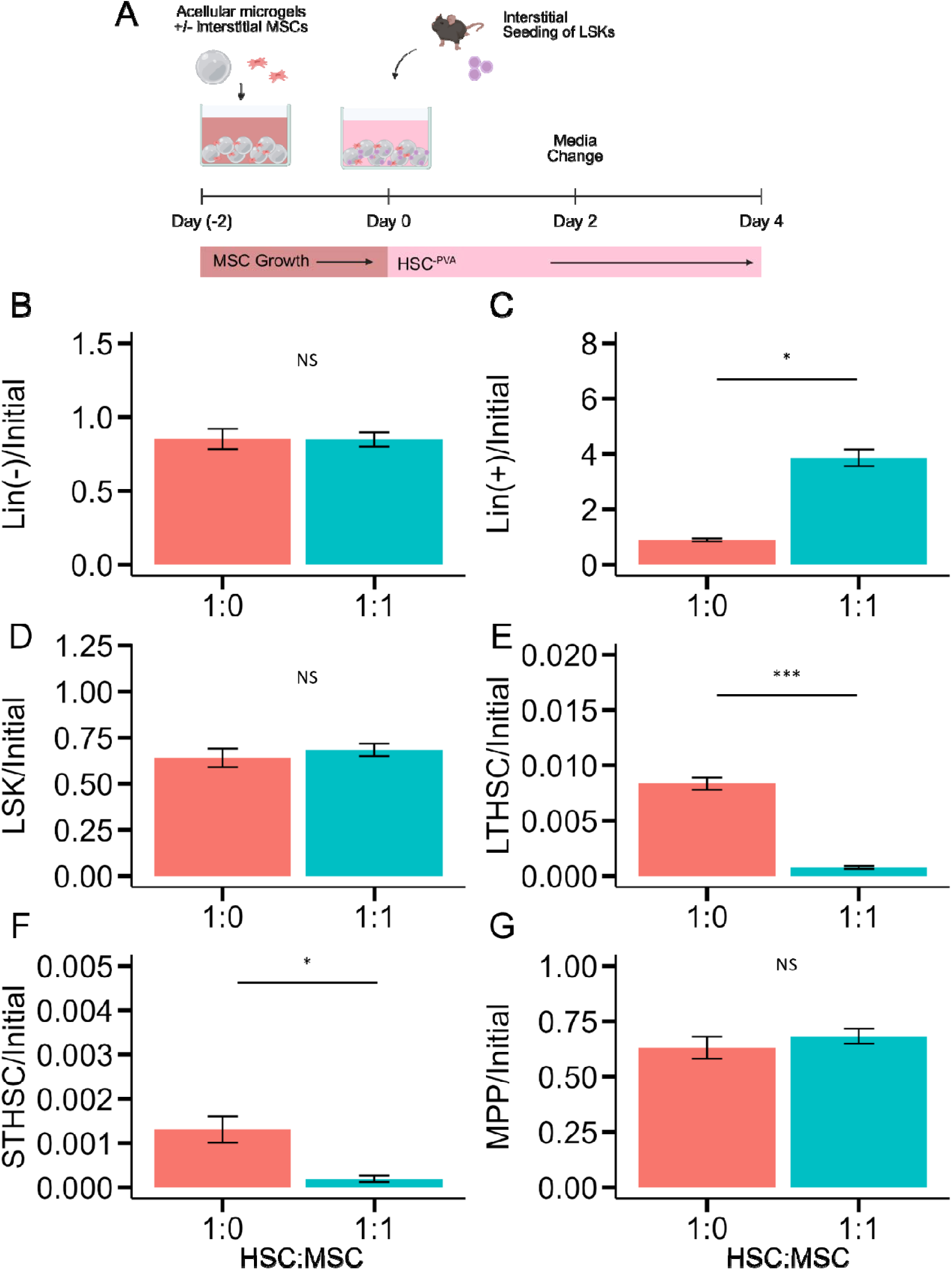
**A** HSCs were interstitially seeded in loosely packed beds of either acellular or MSC-laden microgels and cultured for 4 days prior to flow cytometry analysis. MSCs were seeded interstitially in the bed of microgels. Plots present **B** Lin^-^ cells normalized to the initial seeded number of cells, and **C** Lin^+^ cells normalized to the initial seeded number of cells. The HSPC cell populations were analyzed and normalized against the initial number of ceded cells as follows: **D** LSKs, **E** LTHSCs, **F** STHSCs, **G** MPPs. * p < 0.05, ** p < 0.01, *** p < 0.001, **** p < 0.0001.

## 4. Discussion

Hydrogels have enjoyed extensive use for cell culture due to their ability to recapitulate physiologically mimetic 2D and 3D microenvironments. Here, we investigated the effects of microgel encapsulation of MSCs upon MSC viability and gene expression. In previous work, we observed a roughly 10-fold increase in MSC number over 7 days of culture [41]. In the present study, viability was high at all measured timepoints, and MSCs encapsulated migrate to and colonize the surface of microgels within about 3 days. These observations are consistent with the behavior of human MSCs in degradable poly(ethylene glycol) microgels studied by Mora-Boza et al. [63]. The expression of many genes associated with ECM remodeling and HSC regulation increased over time in culture out to 7 days. *Cxcl12*, *Kitl, Ptn,* and *Vcam1* encode proteins that retain and promote the maintenance of HSCs within the bone marrow [16, 64–69]. The upregulation of these factors highlights the utility of microgel encapsulation of MSCs to promote HSC stemness. Notably, *Spp1* (which encodes the endosteal niche-associated factor osteopontin) is downregulated over time, suggesting that these microgel-encapsulated MSCs more closely mimic the perivascular niche than the endosteal [70]. Osteopontin has also been shown to enhance quiescence in HSCs [71]; future efforts may benefit from exogenous supplementation or engineering of an artificial niche that promotes *Spp1* expression if proliferation is too high. The increase in *Mmp2* and *Mmp9* expression is consistent with the rapid migration of cells through the nanoporous hydrogel matrix, which necessitates metalloprotease-mediated breakdown. The increase in *Fn1* expression suggests cell-mediated remodeling toward a bone marrow-mimetic environment [72] and has been shown to enhance the maintenance of immature HSPCs by our lab [73] as well as by others [21, 74]. Interestingly, the mode of encapsulation was a significant determinant of gene expression. Microgel-encapsulated MSCs increased in their activity over time, while the gene expression of macrogel-encapsulated MSCs declined. Since the polymer itself was held constant in these cases, these data suggest the importance of hydrogel morphology – namely, that reduced diffusive lengths enhance transport of oxygen, nutrients, wastes, and paracrine signals throughout the culture.

We identified a media formulation for effective co-culture of HSCs and MSCs that supported MSC activity and satisfied the primary objective of minimizing HSC exposure to serum-containing MSC media. Removal of PVA from the conventional HSC medium [21, 44] facilitated MSC culture, albeit with reduced proliferation relative to conventional MSC medium. Even with the removal of PVA, MSCs encapsulated in microgels and cultured entirely in HSC^-PVA^ medium suffered poor viability. Because we observed that in 2D culture the HSC^-PVA^ medium can support MSCs, we hypothesize that microfluidic encapsulation presents stresses [75] to MSCs that are ameliorated by serum proteins or factors present in the MSC medium. This was further supported by the observation that HSC^-PVA^ medium could support microgel-encapsulated MSCs, if MSCs were encapsulated and pre-cultured for 2 days in MSC growth medium.

When an equal number of microgel-encapsulated MSCs were cultured with HSCs, we observed an increase in overall hematopoietic progenitor (LSK) maintenance. Increasing the MSC:HSC ratio from 1:1 to 10:1 reduced the overall number of LSKs maintained. However, the most critical LTHSC population was variably affected by the presence of MSCs, with the weight of the combined evidence showing that microgel-encapsulated MSCs selectively preserve LTHSCs. STHSCs were consistently depleted by the inclusion of MSCs, with MPPs slightly increased (1:1 MSC:HSC), then depleted (1:10 HSC:MSC) in response to increasing fractions of microgel-encapsulated MSCs. The polymer wt% was expected to be a significant factor in the maintenance and differentiation patterns of HSCs in this experiment, as it was in previous bone marrow models from our lab [35]. In a previous study by Gilchrist et al., a stiffer (higher wt%) polymer was found to promote HSC maintenance when HSCs and MSCs were co-cultured in a 1:1 ratio in a GelMA hydrogel. We therefore expected a denser, stiffer matrix to enhance stemness in this co-culture study based on microgels; however, the polymer fraction only mildly increased STHSC retention and was not impactful otherwise. This may largely be due to the role of diffusive transport between cell populations (slowed in large macrogels vs. packed beds of microgels) and the localization of HSCs within a macrogel in prior studies versus in between hydrogel microgels in this study.

Cell encapsulation via droplet microfluidics is challenging. It subjects cells to shear stress that can decrease viability or alter cell behavior [76], and flow-focusing approaches with high throughput are an exception to the norm [42, 77]. Because MSCs appear to gradually migrate to the microgel surface over a period of 3-4 days, we expected that interstitial seeding would produce similar effects to MSC encapsulation while streamlining culture setup by eliminating cellular encapsulation within microgels. Nevertheless, the profound depletion of both LTHSCs and STHSCs when both HSCs and MSCs were interstitially seeded serves as motivation to encapsulate MSCs within microgels rather than simply seeding into a scaffold. Depletion of LTHSCs and STHSCs may have been caused by excessive Notch signaling: Delaney et al. observed that low levels of immobilized Delta-like ligand 1 promoted the generation of human CD34^+^ HSPCs from cord blood *in vitro*, but a high density of this ligand was deleterious for HSPCs [78]. While this study shows that direct cell-cell signaling may have negatively contributed to stem cell maintenance, there is significant opportunity to leverage the flexibility of granular collections of microgels (with distinct cell populations within or between microgels) to explore this phenomenon in more detail.

This study represents a first-generation approach toward microgel-based HSC culture. The commonly cited advantages of microgels and granular approaches include enhanced scalability owing to reduced diffusive lengths, the capability to mix discretely tailored microgel populations, injectability, and porosity. Here, we showed the effects of reduced diffusive lengths on MSC activity by comparing microgel culture to macrogel culture. We observed large differences on the basis of gel morphology/encapsulation strategy, supporting the potency of this approach. Most notably, microgel-encapsulated MSCs provide a robust framework to create mosaic models of HSC-MSC crosstalk within the bone marrow and demonstrate promising increases in phenotypic LSKs and LTHSCs.

## 5. Conclusions

We established an approach toward microgel-based, multicellular culture of HSCs with bone marrow-derived MSCs. We found that the encapsulation strategy – specifically, the choice to encapsulate within microgels as opposed within macrogels – can be a significant factor in the expression of physiologically relevant genes associated with matrix remodeling and maintenance of HSCs. We found that the mode of MSC seeding – encapsulated within microgels or interstitially in a bed of microgels – altered the maintenance of interstitially seeded HSCs in this model, motivating the pursuit of more challenging, but biologically crucial, encapsulation of MSCs within microgels for HSC support. The rich landscape of granular hydrogel design motifs presents a diverse array of future design approaches and modifications to tune this artificial bone marrow.

## Acknowledgements

This work was funded by the National Institute of Diabetes and Digestive and Kidney Diseases of the National Institutes of Health, Award Number 2 R01 DK099528. The content is solely the responsibility of the authors and does not necessarily represent the official views of the NIH. The authors are also grateful for further funding provided by the Department of Chemical and Biomolecular Engineering and the Carl R. Woese Institute for Genomic Biology at the University of Illinois Urbana-Champaign. The authors would also like to thank the Functional Genomics Core and the Cytometry and Microscopy to Omics Cores within the Roy J. Carver Biotechnology Center at the University of Illinois Urbana-Champaign. Specifically, we would like to thank Tatsiana Akraiko for assistance with gene expression, and Dr. Marcin Wozniak for assistance with cell sorting.

## Contributions (CRediT: Contributor Roles Taxonomy [79, 80])

**G.B. Thompson:** Conceptualization, Data curation, Formal Analysis, Visualization, Investigation, Methodology, Writing – original draft, Writing – review & editing. **K.M. Kuo:** Investigation, Methodology. **A.J. García**: Supervision, Writing – review & editing. **B.A.C. Harley:** Conceptualization, Resources, Project administration, Funding acquisition, Supervision, Writing – review & editing.

## Data Availability Statement

Data for this work is available upon reasonable request to the corresponding author.

