## Supplementary Information for "A microgel bone marrow model of mesenchymal stem cell paracrine signaling supporting hematopoietic stem cell retention"

^2^ Dept. Materials Science and Engineering

^3^ Cancer Center at Illinois

^4^ Carl R. Woese Institute for Genomic Biology

University of Illinois at Urbana-Champaign

Urbana, IL 61801

^5^ Parker H. Petit Institute for Bioengineering & Bioscience

^6^ George Woodruff School of Mechanical Engineering

Georgia Institute of Technology

Atlanta, GA 30332

**Corresponding Author:**

B.A.C. Harley

Dept. of Chemical and Biomolecular Engineering

Cancer Center at Illinois

Carl R. Woese Institute for Genomic Biology

University of Illinois at Urbana-Champaign

110 Roger Adams Laboratory

600 S. Mathews Ave.

Urbana, IL 61801

**Table S1** Genes and associated Taqman Assay ID numbers for probes used in gene expression experiments.

| **Gene name** | **Taqman Assay ID** | **Reference** |
| --- | --- | --- |
| *MMP2* | Mm00439498_m1 | [35] |
| *MMP9* | Mm00442991_m1 | [35] |
| *Timp1* | Mm01341361_m1 | [35] |
| *Timp2* | Mm00441825_m1 | [35] |
| *Timp3* | Mm00441826_m1 | [35] |
| *Col1a1* | Mm00801666_g1 | [35], [46] |
| *Col4a1* | Mm01210125_m1 | [46] |
| *Fn1* | Mm01256744_m1 | [46] |
| *Tnc* | Mm00495662_m1 | [47], [48] |
| *Lama1* | Mm01226102_m1 | [46], [49] |
| *Mmp13* | Mm00439491_m1 | [50] |
| *Cxcl12* | Mm00445553_m1 | [10] |
| *Vcam1* | Mm01320970_m1 | [51] |
| *Spp1* | Mm00436767_m1 | [51] |
| *Ptn* | Mm01132688_m1 | [10] |
| *Kitl* | Mm00442972_m1 | [10] |
| *Gapdh* | Mm99999915_g1 | [35] |

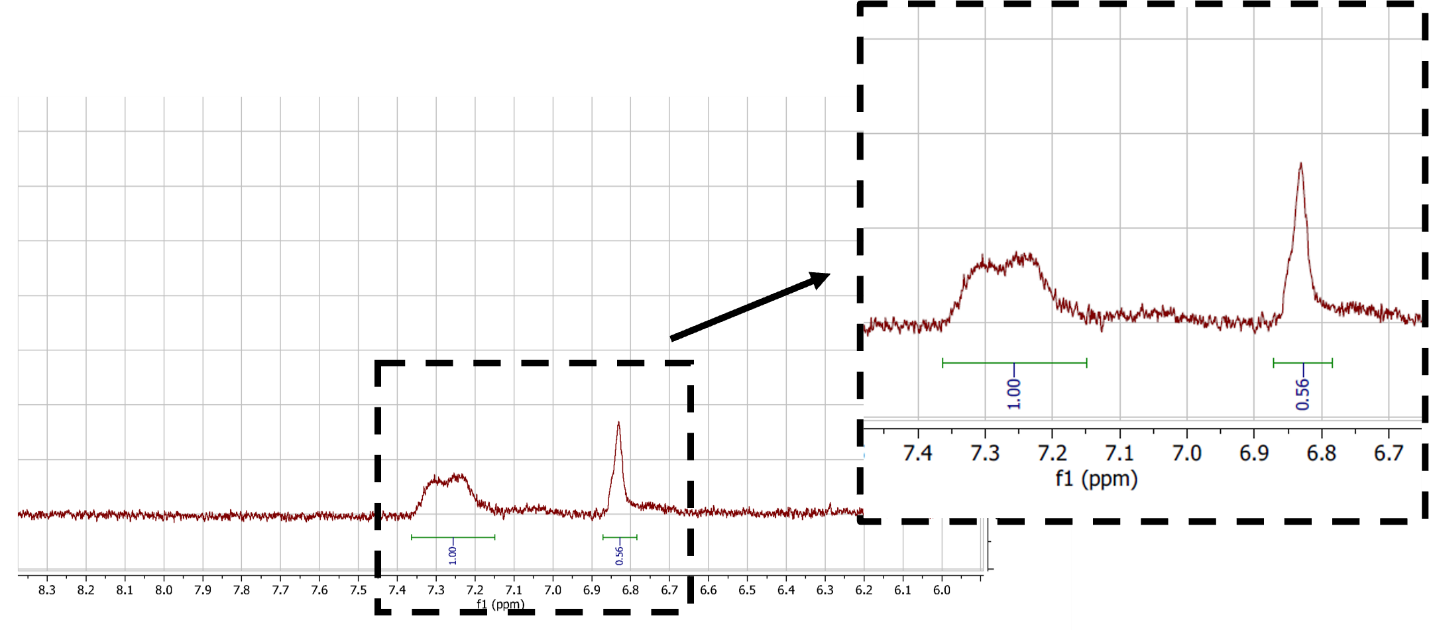

**Figure S1** 1H-NMR spectrum of GelMAL. Figure subsection shows the phenylalanine peak (left) and maleimide peak (right), used for relative quantification of the degree of functionalization.

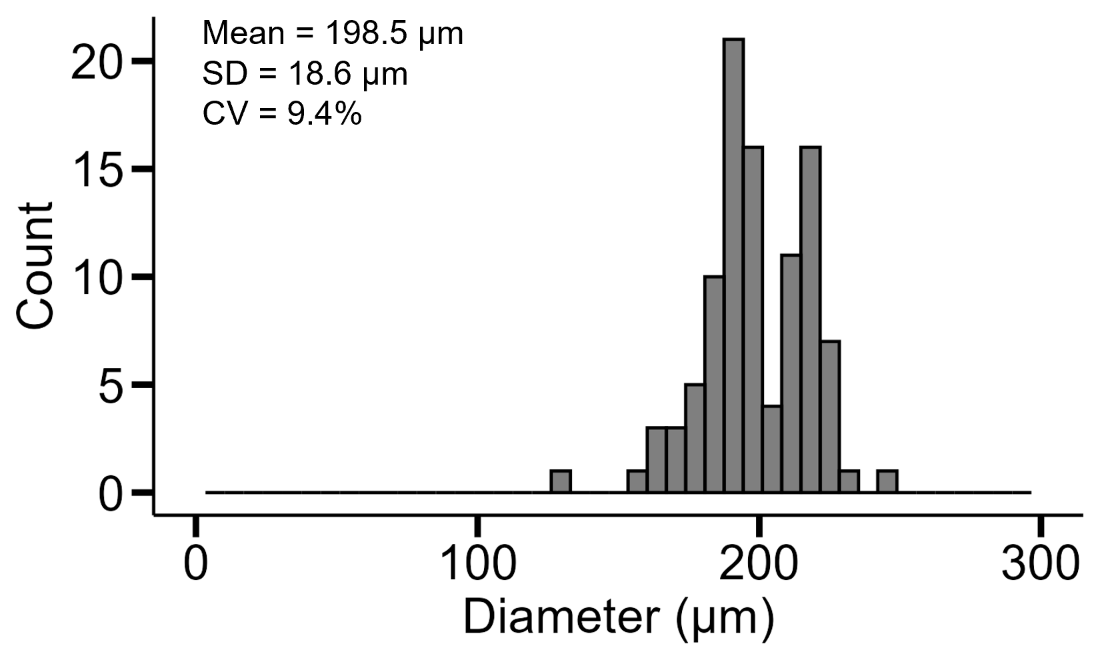

**Figure S2** Representative distribution of 4wt% GelMAL microgel diameter. n = 100 microgels.

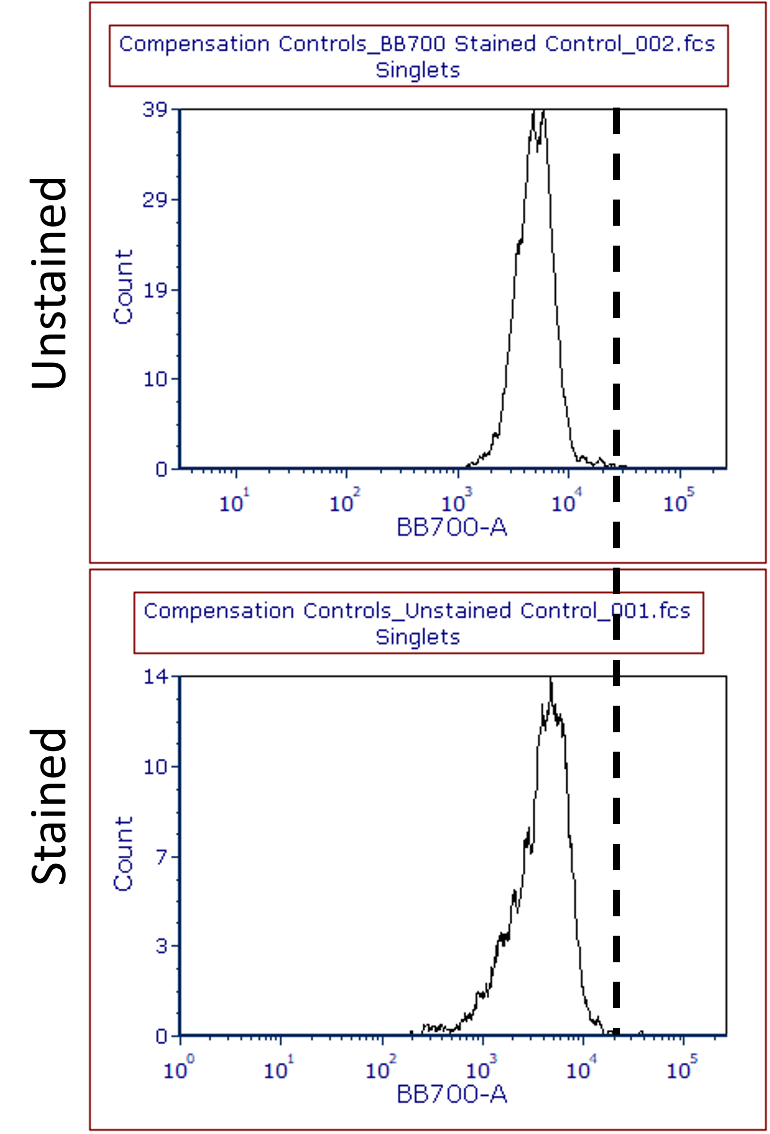

**Figure S3** MSCs were cultured and analyzed for expression of Lineage markers used for HSC analysis. Stained cells fell below the threshold for Lineage marker positivity.

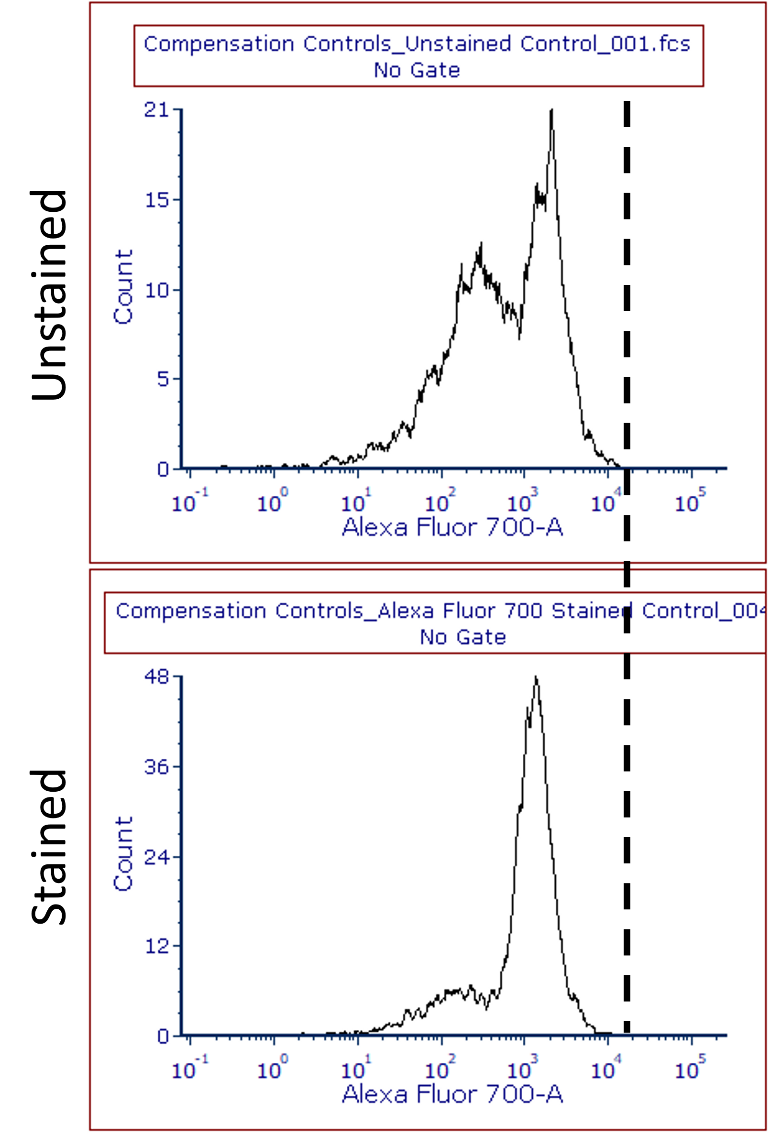

**Figure S4** MSCs were cultured and analyzed for expression of the c-Kit marker used for HSC analysis. Stained cells fell below the threshold for Lineage marker positivity.

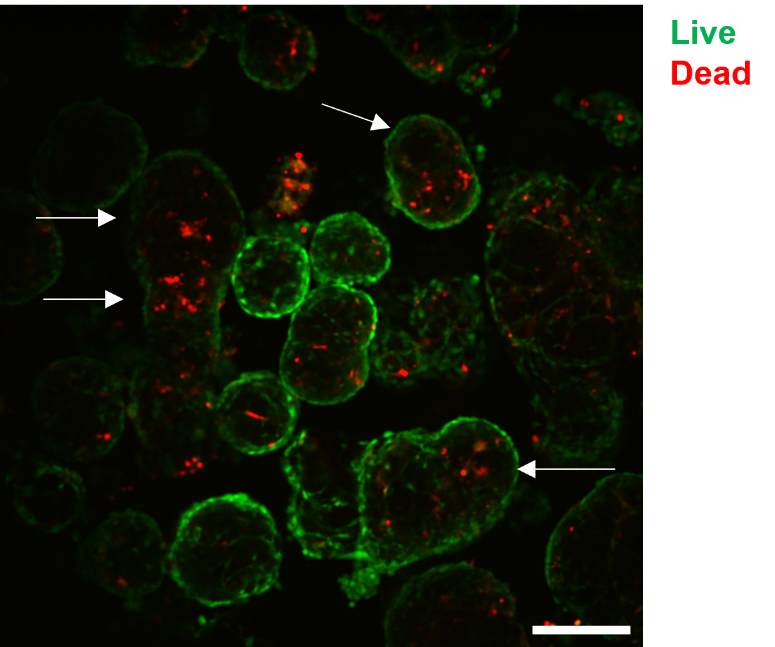

**Figure S5** Representative slice of a z-stack of MSCs grown in 4 wt% GelMAL microgels for 7 days. Dead cells, stained red, appear more frequently at the interior of the microgels. Scale bar 200 µm.

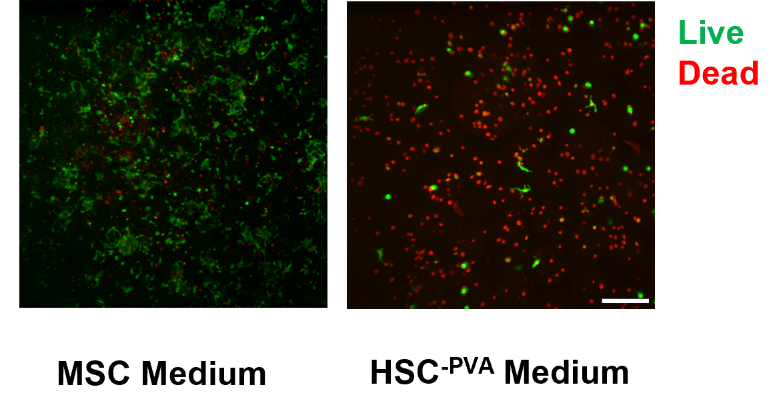

**Figure S6** Representative images of MSCs grown in 4 wt% GelMAL microgels for 4 days. Cells seeded in microgels and cultured in MSC growth medium maintained viability and proliferated, while HSC^-PVA^ medium alone failed to support encapsulated MSCs when. Scale bar 200 µm.

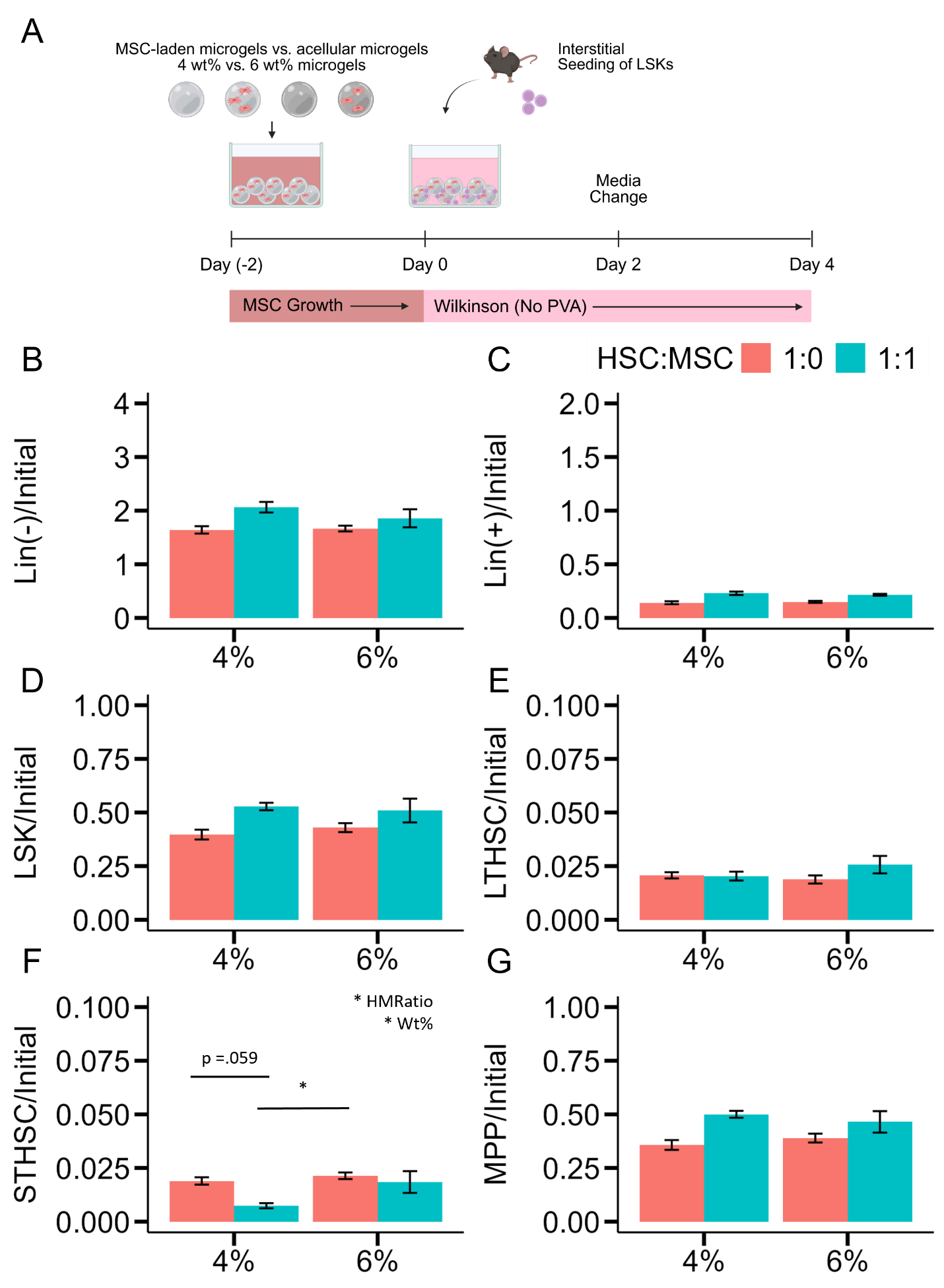

**Figure S7 A** HSCs were interstitially seeded in loosely packed beds of either acellular or MSC-laden microgels and cultured for 4 days prior to flow cytometry analysis. MSCs were encapsulated within microgels comprising either 4wt% or 6 wt% GelMAL. Cells were cultured for 4 days prior to flow cytometry analysis. MSCs were encapsulated within microgels. Plots present **B** Lin^-^ cells normalized to the initial seeded number of cells, and **C** Lin^+^ cells normalized to the initial seeded number of cells. The hematopoietic stem and progenitor cell populations were analyzed and normalized against the initial number of ceded cells as follows: **D** LSKs, **E** LTHSCs, **F** STHSCs, **G** MPPs. The only instance in which GelMAL wt% was a significant factor was for STHSCs. The significance of the factors is present in the top-right of the plot. * p < 0.05.
